# The dynamics of subunit rotation in a eukaryotic ribosome

**DOI:** 10.1101/2021.04.19.440545

**Authors:** Frederico Campos Freitas, Gabriele Fuchs, Ronaldo Junio de Oliveira, Paul Charles Whitford

## Abstract

Protein synthesis by the ribosome is coordinated by an intricate series of large-scale conformational rearrangements. Structural studies can provide information about long-lived states, however biological kinetics are controlled by the intervening free-energy barriers. While there has been progress describing the energy landscapes of bacterial ribosomes, very little is known about the energetics of large-scale rearrangements in eukaryotic systems. To address this topic, we constructed an all-atom model with simplified energetics and performed simulations of subunit rotation in the yeast ribosome. In these simulations, the small subunit (SSU; ~1MDa) undergoes spontaneous and reversible rotations (~ 8°). By enabling the simulation of this rearrangement under equilibrium conditions, these calculations provide initial insights into the molecular factors that control dynamics in eukaryotic ribosomes. Through this, we are able to identify specific inter-subunit interactions that have a pronounced influence on the rate-limiting free-energy barrier. We also show that, as a result of changes in molecular flexibility, the thermodynamic balance between the rotated and unrotated states is temperature-dependent. This effect may be interpreted in terms of differential molecular flexibility within the rotated and unrotated states. Together, these calculations provide a foundation, upon which the field may begin to dissect the energetics of these complex molecular machines.

## 1 Introduction

The ribosome is a massive molecular assembly that undergoes a wide range conformational rearrangements in order to accurately synthesize proteins [1–4]. The precise composition of a ribosome is organism-specific, though it is generally composed of two large RNA (rRNA) chains (~1000-4000 nucleotides in length, each), in addition to a variable number of smaller RNA and protein molecules (~50-100, in total). The overall architecture of the ribosome is commonly described in terms of the large subunit (LSU) and small subunit (SSU), where each has three distinct tRNA binding sites (Fig. 1). During protein synthesis, aminoacyl-transfer RNA (aa-tRNA) molecules must decode the messenger RNA (mRNA). Upon recognition of an mRNA codon, the incoming aa-tRNA molecule binds the ribosomal A site. The nascent protein chain that is attached to the P-site tRNA is then passed to the A-site tRNA through the formation of a peptide bond. After peptide bond formation, the A-site and P-site tRNA molecules are displaced to the P and E sites, respectively. This process, called translocation, leads to a vacant A site, which allows the ribosome to decode the next mRNA frame.

**Figure 1.**
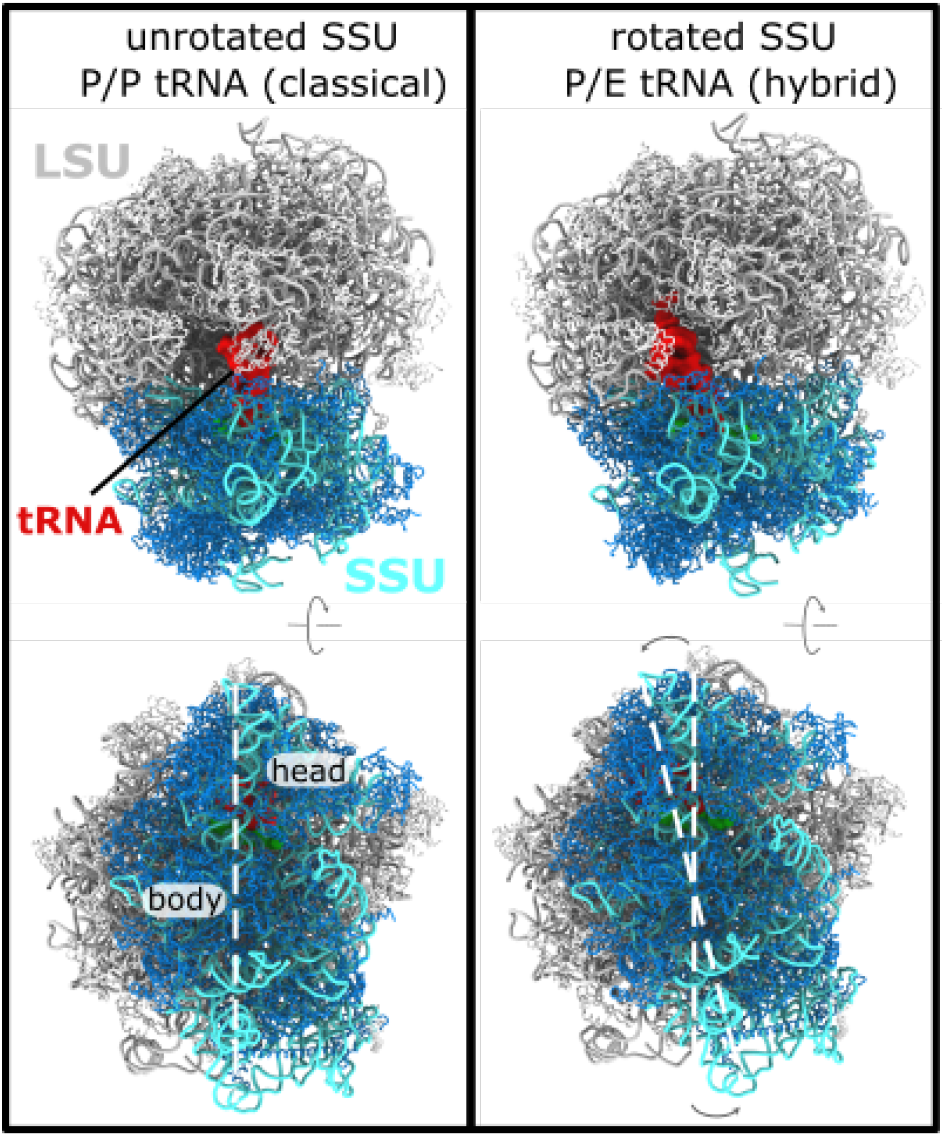
Subunit rotation in a eukaryotic ribosome. The elongation cycle of the ribosome involves numerous large-scale conformational rearrangements. A) In the ribosome, the nascent protein chain is attached to the P-site tRNA molecule (red). A “classical” P/P configuration is shown. In this state, the small subunit (SSU; rRNA:cyan, protein:blue) is described as being in an “unrotated” conformation. Perspective shown in the bottom panel is rotated ~ 90° about the horizontal axis. B) After peptide bond formation, where the nascent chain is transferred to an incoming tRNA molecule, the P-site tRNA adopts a hybrid P/E conformation, where it is displaced toward the E site of the large subunit. P/E formation is also accompanied by a ~ 8° counterclockwise rotation of the SSU, relative to the large subunit (LSU; rRNA:gray, proteins:white). White dashed lines are shown to highlight the relative rotation of the SSU. In the current study, we apply molecular dynamics simulations to probe the dynamics and energetics of SSU rotation in yeast. Structures shown are PDB entries 3J78 (unrotated, classical) and 3J77 (P/E, rotated) [14].

The process of tRNA translocation involves large-scale displacements of tRNA molecules (~40Å), which are facilitated by an elaborate sequence of collective rearrangements within the ribosome. In bacteria, countless cryo-EM and crystallographic structures have been resolved in which the SSU is rotated relative to the LSU (i.e. body rotation [5,6]), or the SSU head is rotated relative to the SSU body (head swivel [7,8] and tilting [9]). To complement these structural snapshots, biochemical measurements [10] and single-molecule studies [11–13] have identified coupling between global SSU rearrangements and tRNA dynamics. Similarly, structural models of eukaryotic [14, 15] and mitochondrial [16] ribosomes have revealed a broad range of orientations that are accessible to the SSU. In addition to rotary-like rearrangements, cryo-EM structures have also revealed that the SSU body may undergo tilt-like rearrangements in eukaryotic systems (called “rolling” [15]). These more recent insights into eukaryotic structure raise new questions into the relationship between SSU motion and tRNA dynamics. For example, how do differences in eukaryotic and bacterial ribosome structure give rise to differential dynamics? What is the relationship between SSU rotation and eukaryotic-specific tilting/rolling? While there is significant interest in understanding these dynamic properties, the biophysical features that govern eukaryotic translation are largely unexplored.

For nearly two decades, advances in structure determination have fueled the development and application of theoretical models to study subunit rotation in bacteria. In the earliest theoretical efforts, coarse-grained models were utilized to perform normal mode analysis [17] and principal component analysis [18], which illustrated how the architecture of the ribosome predisposes it to rotation-like fluctuations of the SSU. Later, highly-detailed explicit-solvent simulations (100ns-1μs) were applied to study smallscale structural fluctuations [19,20], which reinforced the predicted energetic accessibility of rotary-like fluctuations. Explicit-solvent simulations have also been applied to characterize how SSU-LSU bridge interactions may facilitate large-scale rotation [21]. Complementary to explicit-solvent methods, recently-developed coarse-grained models have allowed for spontaneous rotation events to be simulated [22]. In those models, it has been shown how specific subunit bridges can “hand off” the SSU during rotation. While each effort has provided insights into distinct aspects of rotation in bacterial ribosomes, simulations of spontaneous and reversible SSU rotation in a eukaryotic ribosome have not been reported previously.

In the current study, we provide a physicochemical foundation for understanding the dynamics of SSU rotation in eukaryotic ribosomes. Specifically, we developed and applied an all-atom structurebased (SMOG) model [23, 24] to study the dynamics of subunit rotation in the yeast ribosome. This model includes all non-hydrogen atoms (206k atoms, in total), and the energetics are defined to explicitly stabilize the rotated and unrotated conformations. Using this simplified model, we were able to simulate 25 reversible rotation events, where the SSU spontaneously rotated/back-rotated by ~ 8° degrees. With this data set, we provide an initial description of the free-energy landscape associated with SSU rotation. This analysis reveals a distinct sequence of rearrangements at the SSU-LSU interface, as well as a pronounced temperature dependence of the free-energy landscape. Together, these calculations establish a technical and conceptual framework for rigorously characterizing eukaryotic ribosome dynamics, both through theoretical and experimental approaches.

## 2 Materials and Methods

### 2.1 Multi-basin structure-based model

In the current study, we developed a multi-basin structure-based model, where knowledge of the rotated and unrotated conformations (PDB: 3J77 and 3J78 [14]) were used to define the potential energy function. For this, we applied a SMOG-AMBER variant [25] of the SMOG class of structure-based models [24], where the potential energy is given by:

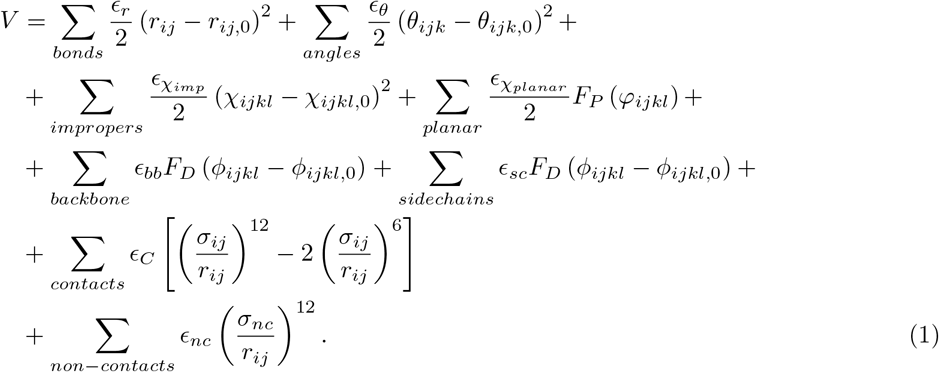

Here, 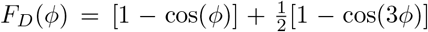 and *F_P_*(*φ_i_*) = [1 – cos(2*φ_i_*)]. The bonded parameters (*r*_*ij*,0_ and *θ*_*ijk*,0_) were obtained from the AMBER03 force field [26]. The planar ring dihedrals were maintained by cosine potentials of periodicity 2. The position of the minimum of each dihedral angle *ϕ*_*ijkl*,0_ was defined as the mean value adopted in the rotated and unrotated structures. This ensures that the dihedral energies in the two structures are isoenergetic. Combined with the AMBER bonded geometry, these terms ensure there is no intrasubunit bias toward either endpoint configuration. Dihedral energies were assigned as described previously [24]. For completeness, we will summarize the details, here. To define the dihedral interaction weights (*ϵ_bb_* and *ϵ_sc_*), dihedrals were first grouped based on the composition of the middle bond. Each dihedral group was given a summed weight of *ϵ_bb_* or *ϵ_sc_*. The ratio 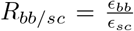 was set to 1 (nucleic acids) or 2 (proteins). *ϵ_bb_* was defined to be equal for protein and nucleic acids. Contact and dihedral strengths were scaled, such that

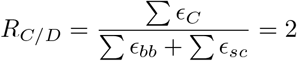

and

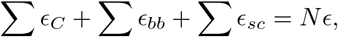

where *N* is the number of atoms in the system and e is the reduced energy unit, which is equal to 2*k_B_T*. Contact pairs were defined using the Shadow Contact Map algorithm with default values [27]. *σ_ij_* was set to 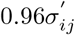 where 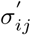 is the interatomic distance between atoms that are in contact in the rotated (or unrotated) configuration. This scaling of contacts was introduced to avoid the artificial expansion of the ribosome that can arise from configurational entropy [25, 28].

To construct the multi-basin force field, the inter-subunit contact pairs found in both structures were combined. The position of the minimum was defined for each atom pair, such that the contacts are isoenergetic (with value *ϵ*_iso_), with respect to the rotated and unrotated conformations (Fig. S1). If *ϵ*_iso_/*ϵ_C_* > 1/2, the contacting pair of atoms was classified as a “common” contact. Applying this criterion, all inter-subunit pairs whose distances are similar in both structures were assigned an isoenergetic distance. All intra-subunit contact pairs were also considered common contacts. Consistent with the dihedral parameters, this isoenergetic assignment of contacts ensures that common interactions do not favor either rotation state. The weights of common contacts (inter-subunit and intra-subunit) followed the scaling rules described above [24]. For the remaining interface contacts (i.e. unique contacts), *σ_ij_* was assigned the value found in the conformation for which the contact is defined: (un)rotated contacts are given the distances found in the (un)rotated conformation. Contacts unique to the rotated configuration were then given an energetic weight of 0.21 and contacts unique to the unrotated configuration were given weights 0.19. An initial parameter sweep was performed to identify weights for which the rotated and unrotated conformations represent pronounced free-energy minima of comparable depths.

Since the available cryo-EM reconstructions resolved different numbers of atoms, minor components of the structure had to be structurally modeled. Consistent with previous efforts [28], position-restraint based techniques were applied to model the missing regions, where the completed region in the alternate structure was aligned to the missing region. With this approach, residues 65-135 of eL24 were reconstructed in the unrotated structure (3J78), and residues 2061-2075 of the 25S rRNA were reconstructed in the rotated structure. After these steps, both models contained 206389 non-hydrogen atoms.

### 2.2 Simulation details

The simulations were performed using the GROMACS software package (v5.1.4) [29,30]. Force field files were generated using SMOG v2.3 [24] with the `SBM_AA-amber-bonds` force field templates [25], which are available through the smog-server.org force field repository. The contacts and dihedrals were then subsequently modified by custom scripts. Two simulations were initiated from the unrotated structure (PDB: 3J78 [14]). To ensure robustness of the results, an additional simulation was initiated from the rotated configuration (PDB: 3J77 [14]). The temperature was maintained via Langevin Dynamics protocols, with a value of 60 in GROMACS units, or a reduced value of 0.49887, for all simulations. Each simulation was continued for at least 7 o 10^9^ time steps. Using the estimate of Yang *et al*. [31], each simulation represents an effective timescale of approximately 15 milliseconds. To allow for equilibration, the first 10^7^ time steps were excluded from analysis. The time step was 0.002 reduced units and configurations were saved every 5000 time steps.

### 2.3 SSU rotation measures

To describe the orientation of the SSU, we extended the definition of rotation angles that were originally introduced for bacterial ribosomes [28,32]. Consistent with previous descriptions [28], the overall strategy is to identify sets of residues that are structurally conserved between the rRNA of the LSU and SSU body in *E. coli* and yeast. Using these structurally-conserved residues, a reference *E. coli* structure is separately aligned to the LSU and SSU body of the yeast ribosome. This allows one to approximate the position of the SSU body and head in terms of rigid body rotations, relative to *E. coli*. The reference *E. coli* model is PDB entry 4V6D [6], where the axis of pure rotation is defined by the structures of the classical and rotated body. With this definition, we decompose the rotation in terms of Euler angles (*ϕ, θ, ψ*). Here, we report the net body rotation as *ϕ*_body_ = *ϕ* + *ψ*, body tilting/rolling as *θ*_body_ = *θ* and the tilt direction as *ψ*_body_ = *ϕ* + *C*. *C* is an arbitrary constant that ensures *ψ*_body_ = 0 reflects tilting motions that are roughly about the long axis of helix 44 in the SSU body.

In the current study, we extended our protocol for defining angles, in order to apply Euler Angle decomposition to the yeast ribosome, while still allowing for direct comparisons with SSU orientations in other organisms. First, we performed STAMP alignment [33] to determine the corresponding *E. coli* numbering of the yeast rRNA residues. STAMP was applied separately to the LSU and SSU body. Then, for the LSU and SSU body, we applied a second, more stringent, condition to define which residues in the rotated yeast structure are structurally conserved with the reference *E. coli* structure. Specifically, we performed least-squares alignment of the STAMP-aligned residues and calculated the spatial deviation of each P atom. Any P atom that deviated by more than 1^Å^A was then excluded in a second round of fitting. After the second round of alignment, all P atoms (including those that were not used for fitting) that are within (above) 1A are included in (excluded from) the fitting group. This process was repeated until the set of included residues converged. These sets of residues, which we will call the “core” groups, were then used for all subsequent angle calculations in the study. All angle calculations were performed using in-house scripts written for use with VMD [34].

## 3 Results

### 3.1 Simulating spontaneous subunit rotation events in a complete ribosome

To study the dynamics of subunit rotation for a eukaryotic ribosome, we applied molecular dynamics simulations with an all-atom model (206,389 atoms) that employs a simplified energetic representation. Specifically, we used a multi-basin structure-based model, which is inspired by similar models for the study of multi-domain proteins [35]. In this model, all non-hydrogen atoms are represented, and the stabilizing energetic interactions are explicitly defined to stabilize the rotated and unrotated configurations of the ribosome. Here, we define the interactions based on cryo-EM structures of a yeast ribosome in the rotated and unrotated conformations (Fig. 1). With this representation, we were able to simulate spontaneous (i.e. without targeting techniques) and reversible transitions between rotated and unrotated conformations (Movie S1).

When interpreting the physical significance of the simulated dynamics, it is important to recognize that the modeled interactions are intended to reflect the effective energetics of the system [36–38]. That is, structures that have been resolved necessarily represent free-energy minima. In the structure-based model, we directly encode these free-energy minima by defining contacts and dihedrals to stabilize the pre-assigned structures. This general approach has been applied recently to study rotation in a bacterial ribosome with a coarse-grained model [22]. In the current study, we extend to an all-atom representation, such that the presented models may later be used to study the precise influence of sterics during tRNA rearrangements in the ribosome. This consideration is motivated by previous simulations of bacterial ribosomes, which have demonstrated the critical influence of molecular sterics on tRNA dynamics during accommodation [39], A/P hybrid formation [40], P/E hybrid formation [41] and tRNA translocation [28]. In order for our model to have future utility to address these motions in eukaryotic ribosomes, it is necessary to employ atomic resolution. Here, we focus on rotation in the absence of a bound tRNA molecule.

In order to characterize the dynamics of SSU rearrangements, one must define appropriate collective coordinates that distinguish between rotated and unrotated orientations of the small subunit (SSU). To this end, we employed Euler Angle decomposition, as used previously to describe bacterial ribosomes [28,32]. Here, this method was generalized (see Methods for details) for use with non-bacterial ribosomes. Consistent with previous efforts to quantify rotation angles in the ribosome [28], we first defined sets of residues within the LSU and SSU body that undergo minimal intra-domain rearrangements during rotation (called the “core” residues). We then aligned reference structures of the core residues to the LSU and SSU body. The orientations of the aligned structures were then quantified in terms of Euler angles, which may be expressed as a net body rotation angle (*ϕ*_body_), a body tilt angle (*θ*_body_) and a corresponding tilt direction (*ψ*_body_)^1^. Pure body rotation leads to *ϕ*_body_ ≠ 0, while *θ*_body_ = 0, where the rotation axis is parallel to that defined by reference structures of *E. coli* [6]. Non-zero values of the tilt angle *θ*_body_ represent any level of deviation from pure rotation. In relation to other studies of eukaryotic ribosomes, the tilt angle measures the so-called “rolling” rearrangement of the SSU body [15]. For consistency with descriptions of SSU head motion [28], we use the term “tilting” to describe such motions. To complete this description of subunit rotation, the direction of tilting is given by *ψ*_body_, where *ψ*_body_ ~ 50° corresponds to the tilting/rolling rearrangement that is apparent in cryo-EM models of the yeast ribosome [14].

Using our structure-based model, we performed several independent simulations, in which a total of 25 rotation and backrotation events were observed (Fig. 2C,D). It is important to note that, while the atomistic resolution imposes significant computational requirements, the simulated events were obtained without the use of enhanced sampling methods, or artificial targeting forces. Rotation events are apparent in the time traces, where there are sharp changes in *ϕ*_body_ of approximately 8° (Fig. 2C). There are also abrupt transitions in the tilting angle *θ*_body_ that coincide with changes in the rotation angle. Thus, this model suggests that subunit rotation and tilting/rolling are not kinetically-separable processes. While one could expect two-state-like behavior in this type of simplified model, it is important to note that it is common for these types of models to reveal sterically-induced free-energy minima [42]. That is, while the models define specific conformations as stable, the imperfect complementarity of steric interactions can impede the motion and lead to long-lived intermediates. However, despite this possibility, we find that the steric composition of the SSU-LSU interface appears to be sufficiently smooth that rotation and tilting can occur simultaneously, and in a two-state manner.

**Figure 2.**
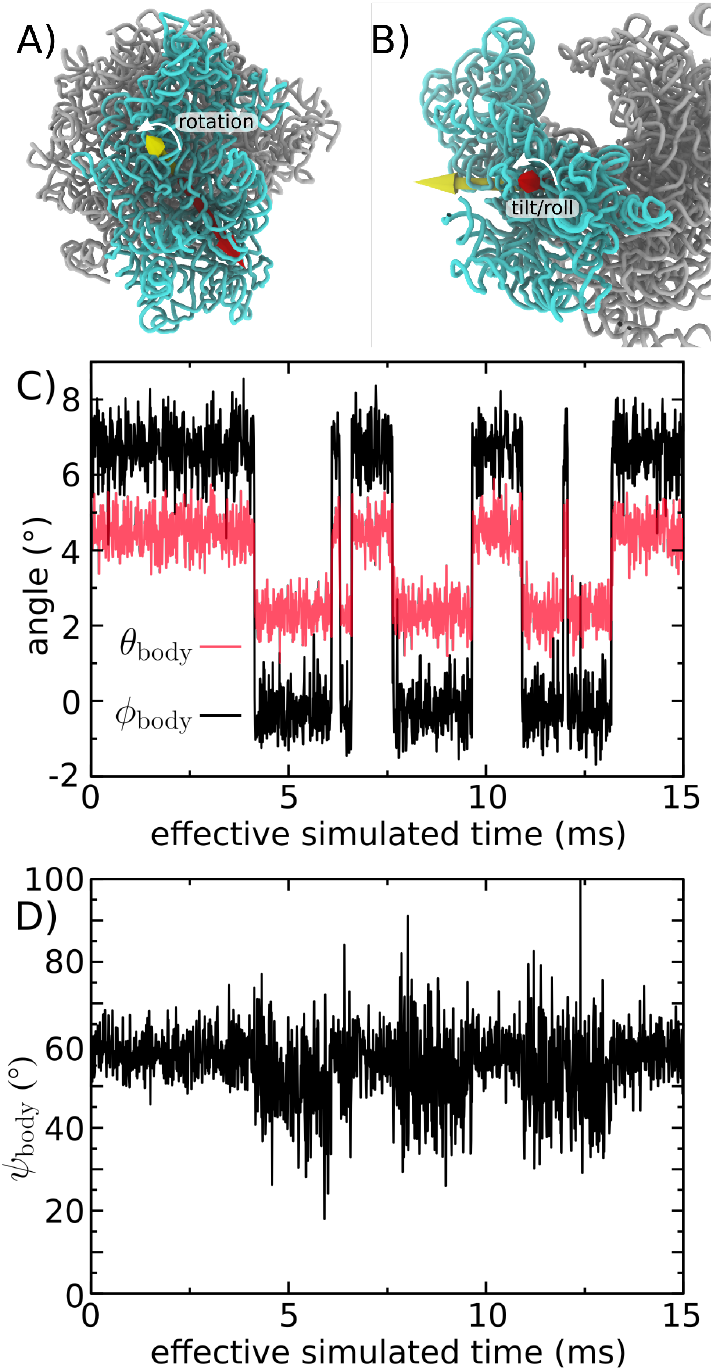
Simulations of subunit rotation in a eukaryotic ribosome. Using an all-atom structure-based model [24], we simulated spontaneous rotation and backrotation of the SSU in yeast. A) Euler angle decomposition was used to describe the orientation of the SSU, where the angle *ϕ*_body_ measures rotation of the SSU body, relative to the LSU. *ϕ*_body_ is defined as rotation about the yellow vector. B) The tile angle θ_body_ describes rotation that is orthogonal to the body rotation angle *ϕ*_body_. The tilt axis (red) may be in any direction perpendicular to the rotation axis (yellow), where the direction of tilting axis is given by *ψ*_body_. In the figure, *ψ*_body_ ~ 60° is shown. C) *ϕ*_body_ and *θ*_body_ shown for a single simulation (of 3, in total). In this model, there are distinct and sharp transitions between the unrotated (*ϕ*_body_ ~ − 1°) and rotated (*ϕ*_body_ ~ 7°) orientations. There are also concommittent changes in *θ*_body_. For reference, the cryo-EM structures [14] correspond to *ϕ*_body_ ~ −1.8° and *θ*_body_ ~ 2.4° for the unrotated state and ~ 8.6° and *θ*_body_ ~ 5.0° for the rotated state. Negative and non-zero angles for the unrotated conformation reflect the relative orientation of the SSU in yeast, relative to the reference bacterial system (*E. coli*). D) In the simulation, the direction of tilting (*ψ*_body_) shifts to slightly higher values as the SSU rotates and tilts. This reveals how the direction of structural fluctuations depends on the global conformation of the ribosome. The effective simulated times are estimated based on previous comparisons with explicit-solvent simulations [31].

With regards to tilting dynamics, we find there are large variations in the tilt direction *ψ*_body_ within the unrotated ensemble (Fig. 2D). These large variations are expected, since the tilt direction is undefined when the tilt angle is zero. Accordingly, when *θ*_body_ is small, structural fluctuations associated with thermal energy can give rise to minimal changes in *θ*_body_ and large changes in *ψ*_body_.

Overall, the presented simulations demonstrate how an all-atom structure-based model can be used to provide a first-order approximation to the dynamics of rotation in a eukaryotic ribosome. Since these simulations describe spontaneous rotation events, the data set provides an opportunity to gain initial insights into the relative timing of inter-subunit contact formation during rotation, as well as the impact of specific interactions on the free-energy landscape.

### 3.2 Quantifying the energy landscape of rotation

A persistent challenge in molecular biophysics is to define low-dimensional measures that can accurately capture the dynamics of complex multi-dimensional processes [43–47]. In addition to posing an intellectual challenge, there is also a practical utility of identifying appropriate coordinates. Specifically, knowledge of appropriate one-dimensional measures can allow one to precisely characterize the relative contributions of individual interactions to biological kinetics. To this end, we explored multiple approaches for describing the simulated SSU rotation events, which together help establish a physical-chemical foundation for the analysis of eukaryotic ribosome dynamics.

Visual inspection of individual simulated time traces suggests there is a strong correlation between SSU rotation and tilting/rolling (Fig. 2C). To better understand the relationship between these motions, we calculated the two-dimensional free energy (−*k_B_T*ln(P)) as a function of *ϕ*_body_ and *θ*_body_ (Fig. 3A). Consistent with the time traces, there are two distinct minima corresponding to the rotated and unrotated ensembles. However, rotation and tilting are not perfectly correlated, and there is a notable degree of variability in *θ*_body_ within each ensemble. In the unrotated ensemble (low *ϕ*_body_ values), the tilt angle samples values that range from 1° to 4°. Similarly, in the rotated ensemble (high *ϕ*_body_ values), *θ*_body_ spans a wide range of values (2.5 – 6.5°). Interestingly, as the body tilts/rolls, the direction of tilting also shifts. Specifically, the direction of tilting (*ψ*_body_) within the unrotated ensemble is typically around 50°, while the rotated ensemble represents configurations for which the tilt direction is towards higher *ψ*_body_ values. This illustrates how, upon reaching the rotated/tilted ensemble, the accessible tilting fluctuations change in character. Based on the current simulations, it is not clear whether this change in tilting dynamics has a specific biological role, though it is possible that it may help coordinate tRNA dynamics during hybrid formation and/or translocation.

**Figure 3.**
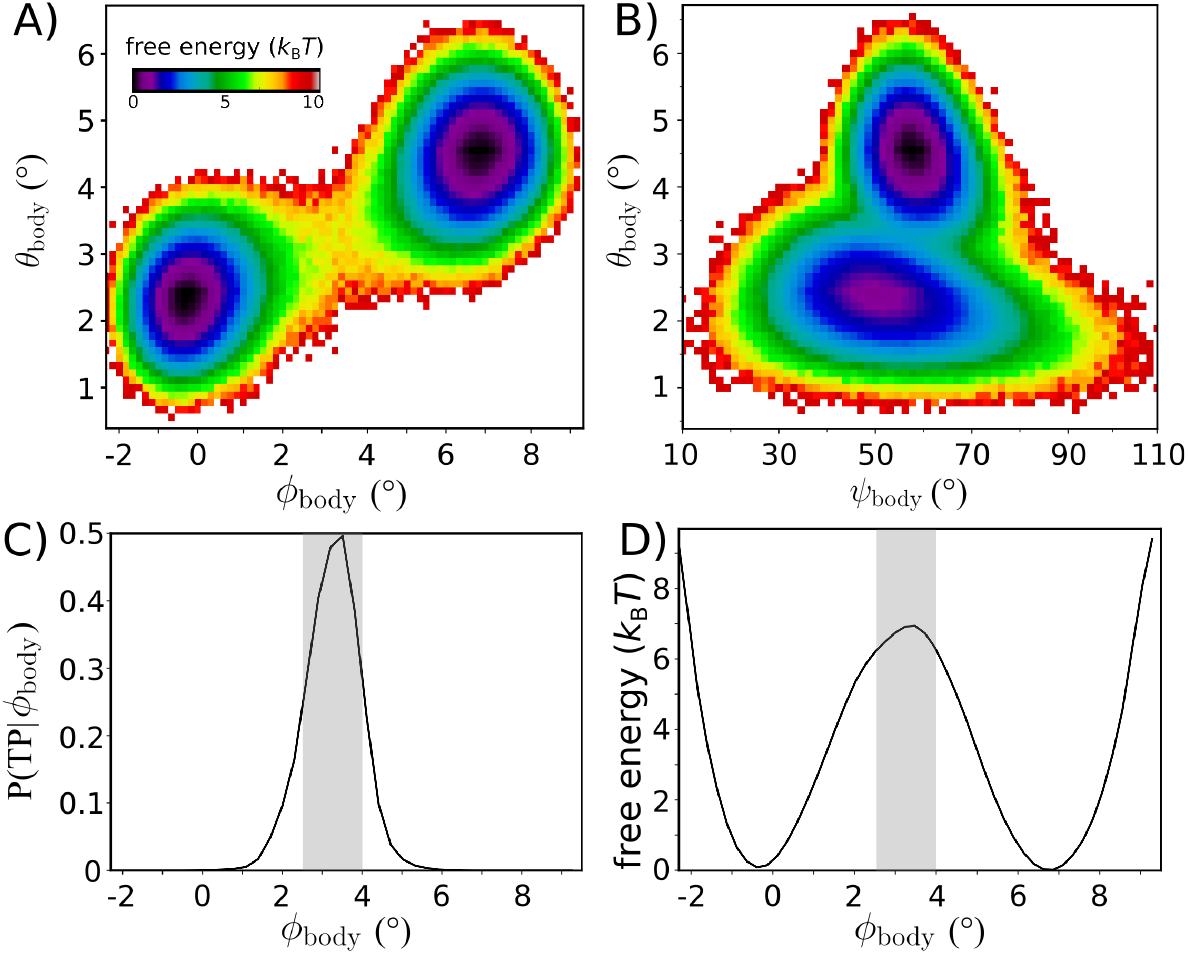
The free-energy landscape of rotation. Equilibrium simulations of the intact yeast ribosome (Fig. 2) were used to characterize the energetics of subunit rotation. A) The free-energy as a function of *ϕ*_body_ and *θ*_body_ suggests the barrier for rotation is ~ 7*k_B_T*. Rotation and tilting are correlated, though the simulations reveal that tilting motions are widely accessible within the unrotated ensemble. When *ϕ*_body_ ~ — 1, the tilting angle *θ*_body_ can adopt values up to ≈ 4°. Similarly, there is a range of ~ 4° in the tilt angle for the rotated ensemble. This indicates that, while rotation and tilting are correlated, the motions are only weakly coupled. B) Free-energy as a function of the tilt angle *θ*_body_ and direction of tilting *ψ*_body_. Comparison of cryo-EM structures would suggest a single direction of tilting, though the simulations indicate that as the ribosome adopts more tilted orientations, the direction of tilting shifts to slightly higher values of *ψ*_body_. As a result, it is appropriate to describe tilting in terms of a twist-like rearrangement that is concomitant with rotation. C) While rotation/tilting involves a complex combination of motions, the kinetics of rotation is described well by the single coordinate *ϕ*_body_. Specifically, the probability of being on a transition path as a function of *ϕ*_body_, *P*(*TP* |*ϕ*_body_), adopts a peak value of ~ 0.5. Gray area indicates the identified TSE region. D) The free-energy as a function of *ϕ*_body_ also yields a barrier that is comparable to that of the two-dimensional landscape shown in panel A.

Since rotation and tilting are found to be correlated, we asked whether a one-dimensional description can be sufficient to quantify the free-energy barrier and kinetics associated with SSU motion. To determine whether the rotation angle *ϕ*_body_ provides a kinetically-meaningful approximation to the free-energy barrier, we employed multiple independent analysis strategies. First, we applied transition path analysis [48,49] to assess whether *ϕ*_body_ can unambiguously identify the transition state ensemble (TSE). That is, we calculated the probability that the system is undergoing a transition (i.e. is on a Transition Path), as a function of *ϕ*_body_: *P*(TP|*ϕ*_body_). If *ϕ*_body_ is able to unambiguously identify the TSE, then *P*(TP|*ϕ*_body_) will adopt the diffusion-limited value of 0.5. Here, we find that P(TP|*ϕ*_body_) reaches a peak value of ≈ 0.5, indicating that *ϕ*_body_ is suitable for describing the underlying barrier. Further, this allows us to identify the position of the TSE: *ϕ*_body_ ~ 3.5°. As a point of comparison, in the study of protein folding [48,49], a coordinate *ρ* is often considered “good” if *P*(TP|*ρ*) reaches values that are greater than ~ 0.4.

In addition to capturing the TSE, we find that *ϕ*_body_ is able to recapitulate the long-time dynamics of SSU rotation. To illustrate this, we used a Bayesian inference approach [49] to calculate the diffusion coefficient as a function of *ϕ*_body_. This produced a nearly-uniform value of the diffusion coefficient *D* that was approximately 0.0025(degrees)^2^/*τ*_ru_. *τ*_ru_ is the reduced time unit. We used this value of *D* to estimate the mean first passage time via the relation:

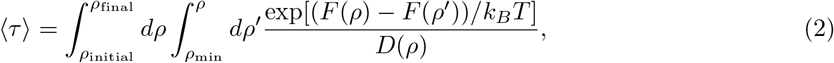

where *F*(*ρ*) is the free-energy as a function of *ρ* = *ϕ*_body_. *ρ*_final_ and *ρ*_initial_ are the values of *ϕ*_body_ at the free-energy minima, and *ρ*_min_ is the smallest accessible value of the coordinate. According to this calculation, the inferred timescale is estimated at 10^6^*τ*_ru_. This is comparable to the apparent timescale (total simulated time divided by the number of observed transitions) of ~ 1.8 × 10^6^*τ*_ru_. Agreement between the apparent and inferred timescales is consistent with *P*(TP|*ϕ*_body_) reaching the diffusionlimited value of 0.5. It is important to note that, even when using intuitively-defined coordinates, there is no guarantee they will be able to accurately recapitulated the long-time kinetics. For example, in a recent study of SSU rotation in bacteria [22], it was shown that a similar coordinate is unable to capture the TSE, and the coordinate dramatically underestimates the height of the free-energy barrier. Fortunately, here, our analysis indicates that *ϕ*_body_ may be used to probe the factors that govern SSU rotation in a eukaryotic ribosome.

### 3.3 Simulations implicate millisecond-scale dynamics of rotation

Since *ϕ*_body_ can be used to precisely describe the free-energy barrier for rotation, we next compared the predicted free-energy barrier with biological kinetics. For this, we use the expression:

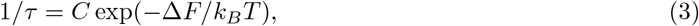

where Δ*F* is the height of the free-energy barrier and *C* is the average barrier-crossing attempt frequency. The free-energy barrier along *ϕ*_body_ is Δ*F* ~ 6 – 7*k_B_T*, and we used a barrier-crossing attempt frequency for rotation *C* that was previously estimated from explicit-solvent simulations [19]. This conversion between the free-energy barrier and kinetics implicates a mean first-passage time *τ* that is in the millisecond regime (1-5 ms). When interpreting this estimate of the timescale, it is important to recall that C was obtained for a bacterial ribosome, which is ~ 70% the mass of the yeast ribosome. In accordance to Stokes-Einstein scaling for rotational diffusion, this would lead one to anticipate that rotational diffusion in yeast will be reduced by a factor of roughly 1.5. Further, since *C* is linearly proportional to the diffusion coefficient [19], the estimated timescale is likely underestimated by a comparable factor. Accordingly, a timescale of 1-5 ms represents a lower-bound on the mean first-passage time for SSU rotation in our model.

Even though the employed model is not intended to provide a complete description of ribosome energetics, the predicted timescale for SSU rotation is generally compatible with the rate of protein synthesis in the cell. The average rate of protein synthesis in yeast has been estimated to be approximately 10 amino acids per second [50], which imposes an upper limit of 100 ms on the timescale of any individual substep. However, there is no evidence that SSU rotation is rate limiting, which would be consistent with rotation kinetics occuring on substantially shorter timescales. In terms of energetics, this empirically-imposed upper bound on the timescale indicates that the barrier height in vivo is necessarily less than ≈ 10*k_B_T* (estimated according to ref. [19]). In terms of modeling considerations, a difference in barrier height that is only a few *k_B_T* can easily be accounted for by minor changes in the energetic representation. For example, increasing the rotated and unrotated contact strength by ~ 2% would increase the barrier by ~ 4*k_B_T*. To make this estimate, we assume that most contacts will primarily stabilize the endpoint conformations. Since there are approximately 500 unique contacts in each endpoint, each of strength ~ 0.2 reduced units (1r.u.= 2*k_B_T*), an increase of 2% would stabilize the endpoints by 4kBT. At the simulated temperature, the reduced energy unit is equal to 2kBT. Since the mechanistic aspects of ribosome dynamics are often robust to parameters changes on this scale [51], we will consider the deviations between the predicted and in vivo upper-bound barrier height to be minimal. Taken together, the compatibility of the timescales for elongation *in vivo* and the simulated rotation events suggests the model represents an appropriate first-order approximation to the energetics of subunit rotation.

### 3.4 Asynchronous dynamics of subunit bridge interactions during rotation

While applying rigid-body descriptions of biomolecular dynamics is often motivated by analogies with macroscopic systems, the scale of thermal energy in molecular systems gives rise to a fundamentally distinct relationship between structure and dynamics. In the cell, solvent introduces energetic fluctuations that are of the same scale (*k_B_T*) as the interactions that maintain structural integrity. This leads to heterogeneous and anisotropic structural fluctuations [52] that can manifest in the form of large-scale global motions [18], as well as more localized distortions and partial unfolding [53,54]. In contrast, a corollary of rigid-body arguments would be that all intersubunit bridge interactions should simultaneously interconvert as the SSU rotates. However, such a process would likely be associated with a large free-energy barriers that would lead to prohibitively slow dynamics.

To better understand how molecular flexibility can facilitate collective rotation of the SSU, we used our simulated transitions to explore the ordering of intersubunit bridge rearrangements. Specifically, we calculated the probability that each rotated and unrotated contact is formed: 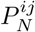. *N* refers to the conformation (rotated, or unrotated) and ij denotes the atom pairs involved in a contact. Here, uniquely rotated/unrotated contacts are defined as atomic interactions that are present (i.e. proximal atoms) in only one of the conformations. Further, to be classified as “unique,” the atom pair must be substantially farther apart (see Methods) in the alternate conformation. We then calculated the average probability of all unique contacts that are defined with a specific atom: 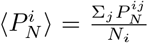, where *N_i_* is the number of contacts that are defined with atom *i*. We then calculated 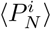 as a function of the rotation angle *ϕ*_body_, for both the rotated (Fig. 4A) and unrotated (Fig. 4B) contacts. Since the contact probabilities will be proportional to the stability imparted by a specific interaction, such analysis allows one to categorize interactions in terms of their contributions to the free-energy barrier and biological kinetics.

**Figure 4.**
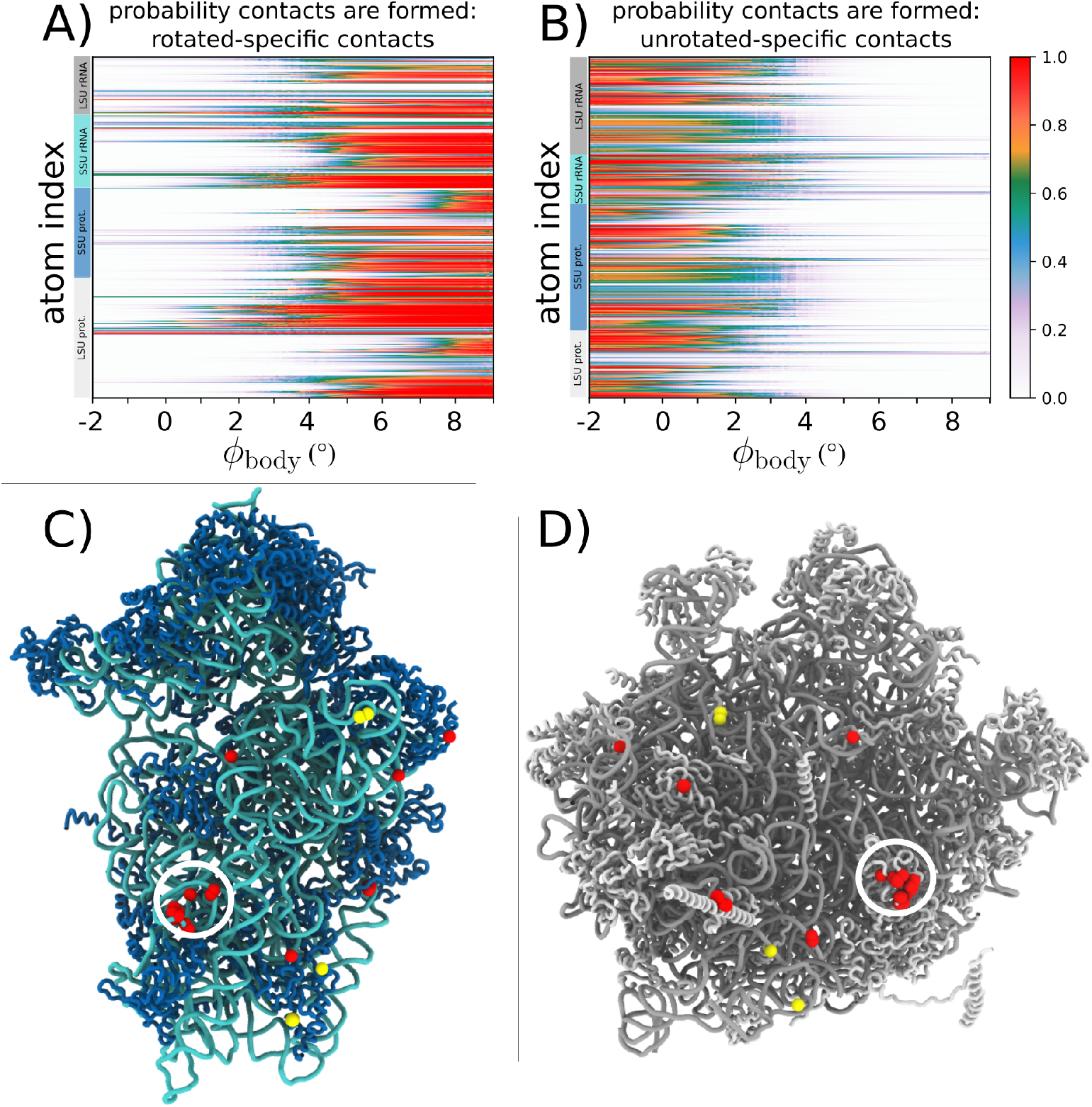
Contact analysis reveals sequential displacements at the SSU-LSU interface. A) Average fraction of rotated-specific inter-subunit contacts formed, by atom, as a function of *ϕ*_body_. Many rotated-specific contacts are formed by the time the ribosome reaches the TSE (*ϕ*_body_ ~ 3.5°). B) In contrast to the rotated-specific contacts (panel A), the unrotated specific contacts are very rarely formed in the TSE. C) Structure of the SSU, viewed from the SSU-LSU interface. Red (yellow) spheres indicate which atoms have at least one rotated-specific (unrotated-specific) contact formed in more than 80% of the TSE frames. D) Structure of the LSU, viewed from the SSU-LSU interface, shown in the same representation as in panel C. While there are only four atoms involved in unrotated contacts in the TSE (yellow), there are clear clusters of rotated contacts formed in the TSE (red). In particular, contacts between h14 and protein L23 (also called uL14, found in bridge B8; circled) are frequently formed, indicating they can have a strong influence on the free-energy barrier and kinetics of rotation.

Contact analysis reveals how specific bridge interactions can have differential effects on the global kinetics of the system. We find that most unrotated contacts break early in the rotation process (Fig. 4B), where almost no interactions are still formed by the time the ribosome reaches the TSE (*ϕ*_body_ ~ 3.5°). This suggests that, while contacts that are unique to the unrotated conformation can contribute to the stability of the classical configuration, these interactions are unlikely to directly affect the free-energy barrier. In contrast, the rotated contacts form over a broad range of *ϕ*_body_ values, where some contacts are likely to form prior to the system reaching the TSE (Fig. 4A).

To visually depict which interface regions are likely to have the strongest influence on the free-energy barrier, we identified all atoms that have at least one contact that is formed 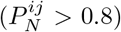 in the TSE. Consistent with Figure 4A/B, there are only four unrotated contacts that are likely to be formed (yellow spheres in Fig. 4C/D). This is in sharp contrast with the dynamics of the rotated contacts, for which there are clusters of formed contacts scattered across the subunit interface. In particular, there is a dense cluster of formed contacts that are centered around bridge B8, which is formed between protein L23 and SSU helix 14. This suggests that, during rotation, the flexibility of the bridge allows it to “reach out” and form rotated contacts before the SSU body has fully transitioned to a rotated configuration. This finding suggests new ways in which experiments may modulate rotation kinetics in eukaryotic ribosomes. For example, it may be possible to introduce mutations to protein L23 that will specifically impact the rotated and TS ensembles, while leaving the stability of the classical configuration unperturbed. Together, these calculations provide a physical framework that can guide the development of next-generation experimental techniques that will be able to control biological dynamics.

### 3.5 Molecular flexibility leads to temperature-induced population shift

In addition to characterizing the transition-state ensemble and the dynamics of SSU-LSU bridge interactions, we next asked whether the balance between rotated and unrotated ensembles is likely to be temperature-dependent. To explore this possibility, we calculated the relative free-energy of the rotated and unrotated ensembles as a function of temperature. We find that increases in temperature are associated with increased stability of the rotated ensemble (Fig. 5A). As described below, one can understand this effect as arising from an increase in mobility of a localized region within the LSU upon rotation.

**Figure 5.**
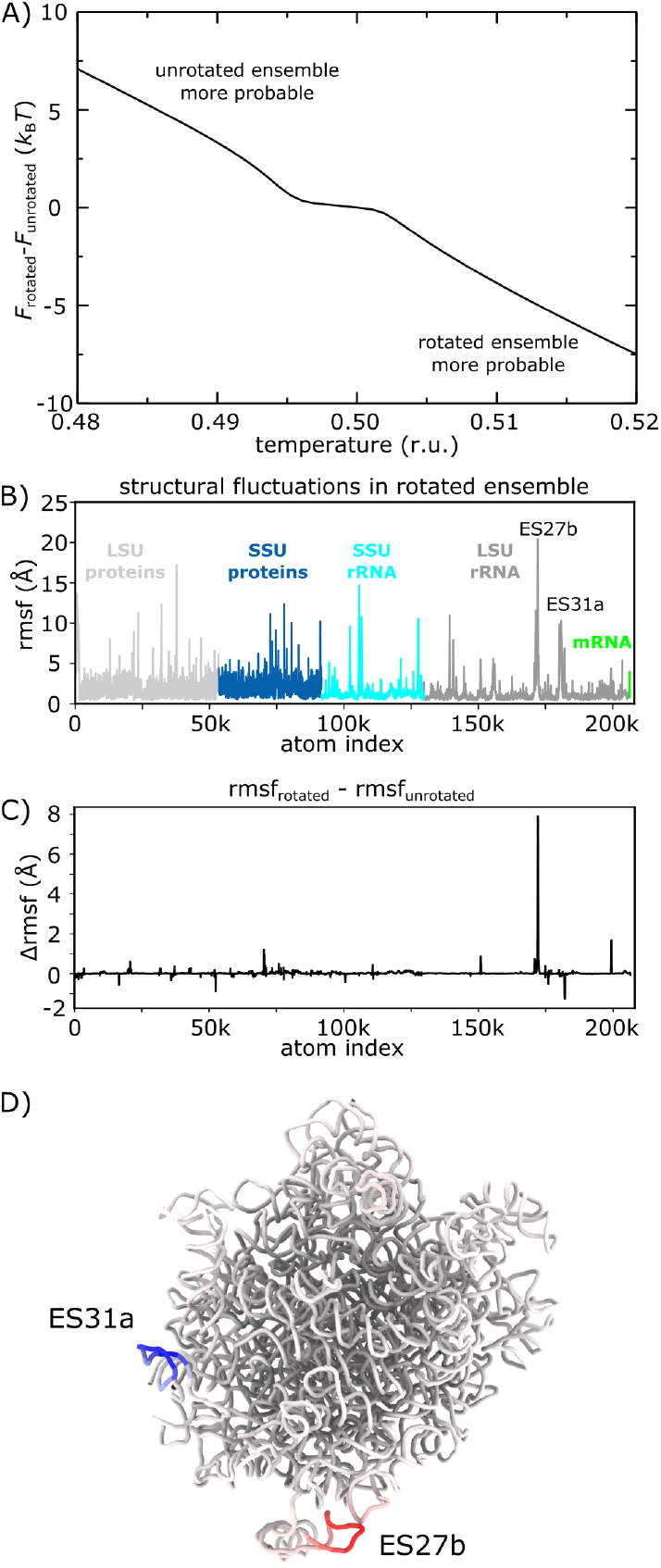
Temperature-dependent free-energy landscape. The presented simulations reveal how the free-energy landscape is influenced by changes in molecular flexibility upon rotation. A) Difference in free-energy of the rotated (5.40° < *ϕ*_body_ < 7.83°) and unrotated (—1.12° < *ϕ*_body_ < 0.82°) ensembles, as a function of temperature. Positive values indicate the free-energy of the rotated ensemble is higher. As T is increased, the free-energy of the rotated ensemble monotonically decreases, relative to the unrotated ensemble. B) Spatial root mean squared fluctuations (rmsf), by atom, for all simulated frames within the rotated ensemble. Peaks in the protein regions are typically due to marginally ordered tails at the peripheral regions of the ribosome. C) Difference between rmsf values calculated for the rotated and unrotated ensembles: Δrmsf. Positive values indicate elevated mobility in the rotated ensemble. Consistent with the temperature-induced shift in population towards the rotated ensemble (panel A), there is one region of the LSU rRNA that is significantly more flexible in the rotated ensemble (expansion segment 27b, ES27b). D) Structural representation of the LSU rRNA, colored by Δrmsf values (blue to red). There is a large increase in flexibility of ES27b, suggesting that rotation allows this region to adopt a wider range of configurations and provide an entropic drive towards the rotated ensemble. This observation is consistent with the lack of electron density obtained for this region in a cryo-EM reconstruction of the rotated state [14].

To probe the temperature dependence of the energy landscape, we pursued a free-energy perturbation (FEP) approach. That is, we calculated the free-energy at arbitrary temperatures according to: *F*(*ϕ*_body_) = −*k_B_T* ln(*P*′(*ϕ*_body_)), where *P*′(*ϕ*_body_) represents a probability distribution for which each simulated frame was assigned a weight:

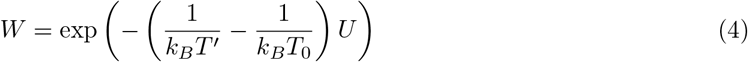

*U* is the potential energy of the simulated snapshot, *T*_0_ is the temperature of the simulation, and *T*′ is the temperature of interest. When *T*_0_ = *T*′ (i.e. original simulated distribution), *W* reduces to 1. Before discussing the results obtained from the FEP calculations, it is important to explain the rationale and assumptions that are implicit in these types of methods. The first consideration was the scale of the simulations. That is, even though simulations with the simplified model are faster (many orders of magnitude) than simulations with explicit-solvent models, the size (>200,000 atoms) and effective timescale (milliseconds) remains very computationally demanding (~ 10^10^ simulated timesteps, per simulation). Thus, it is not practical to obtain equilibrium sampling for a range of temperatures. The second consideration that supported the use of FEP was that the distributions calculated from different simulations were highly reproducible (not shown). This suggests the acquired sampling provides adequate coverage of the local phase space, which is necessary for FEP techniques to provide reliable estimates. The final consideration was that we are interested in temperature effects over a relatively small temperature range. That is, we considered a temperature range of ±2%, where dramatic changes in the accessible conformations of the system are not likely to occur. This is an important point, since one may only extrapolate free energies using perturbation techniques if the perturbed parameter/s (i.e. temperature) would primarily redistribute the probabilities of well-sampled regions of phase space. Since the presented simulations were performed at a temperature for which which only modest changes in flexibility are expected with this model [55], it is reasonable to assume that the accessible range of configurations will not be altered dramatically by small changes in temperature.

We find that the rotated ensemble is favored as temperature is increased (Fig. 5A). For this, the rotated and unrotated ensembles were defined as 5.4 < *ϕ*_body_ < 7.86° and −1.12 < *ϕ*_body_ < 0.82°. The difference in free-energy of the two ensembles was then defined as Δ*F* = *F*_rot_ – *F*_unrot_ = *k_B_T* ln(*P*_unrot_/*P*_rot_), where *F*_N_ is the free-energy of ensemble *N*. For the reported simulations, the free-energy of the two ensembles was comparable (Fig. 3D). However, we find that the relative stability of the rotated ensemble increases with temperature. This suggests that differences in the configurational entropy of the rotated and unrotated ensembles can lead to a distinct temperature-dependence of the distribution between these states. It is important to note that, since the current model does not provide an explicit treatment of the solvent, the predicted trend is due solely to the contributions of the configurational entropy of the ribosome. Based on available data, it is not known whether solvation entropy will provide a significant contribution to this free-energy difference. If future experiments corroborate the observed trend, then one may infer that configurational entropy considerations are sufficient. If an opposite trend were found in experiments, this would suggest that solvation entropy is the dominant contributor. However, since the majority of the SSU-LSU interface is maintained in the rotated and unrotated states, it is reasonable to expect that solvation entropy changes will be minimal.

To provide a structural interpretation for the origins of the temperature-dependent distributions, we considered the flexibility of the ribosome in each ensemble. To assess the differential flexibility of the ribosome, we calculated the spatial root mean-squared fluctuations (rmsf) of each atom, where the average was calculated separately for the unrotated and rotated (Fig. 5C) ensembles. In both cases, the rmsf values are highly heterogeneous, where the stalk regions are most flexible, alongside some peripheral protein tails. To identify which regions of the structure contribute to the observed temperature-dependent dynamics, we calculated the difference in rmsf values Δrmsf = rmsf_rot_–rmsf_unrot_ for each atom (Fig. 5D). This reveals that the differences in flexibility within the ensembles are primarily centered around a small region of H63 (expansion segment 27b, ES27b: red in Fig. 5D). Interestingly, there is also one region in the LSU rRNA involved in the formation of a bridge interaction (ES31a in H79; blue in Fig. 5D) that exhibits a decrease in mobility, though the attenuation of mobility is of a smaller scale than the increase in mobility of ES27b.

In summary, as the ribosome rotates, the ES27b region can dissociate from the SSU interface and adopt a wide range of configurations. This increased mobility can then entropically drive the ribosome towards the rotated ensemble. At a qualitative level, this type of behavior of reminiscent of the notion of an entropic spring in polymer physics, where extension-associated reductions in entropy lead to an effective contractile force. There have also been similar examples in the study of protein conformational rearrangements, where increased mobility has been suggested to act as an entropic counterweight [56]. Accordingly, the current study helps elucidate the interplay between flexibility and dynamics in the ribosome, while revealing a common theme with simpler molecular systems.

## 4 Discussion

While structural methods can reveal atomic details of the ribosome at various stages of function, biological kinetics are controlled by energetics. Due to the complex character of ribosomal motions, directly quantifying these energetics through experiments has proven to be extremely difficult. Thus, there is a need for a quantitative physical foundation, with which precise experimental measurements may be devised and executed. To this end, molecular simulations represent a powerful approach that can help guide the development of next-generation experiments. That is, rather than attempting to predict the exact behavior of a complex assembly, simulations may be used to establish broad trends and relationships. These insights can then suggest experimental strategies that will be able to isolate the factors that control biological dynamics in the cell.

In the presented study, we illustrate how molecular simulations of a eukaryotic ribosome can provide initial insights into the relationship between molecular flexibility and kinetics. For example, we find that rotation is best described in terms of asynchronous motions that are facilitated by heterogeneous flexibility of the ribosome. As another example, we find that the flexibility of a specific helical region (ES27b) can increase upon rotation. This increase in mobility is accompanied by an increase in configurational entropy that can drive the rotation process. In terms of experiments, these predictions suggest that it may be possible to alter the dynamics of large-scale processes by introducing localized modifications to the ribosome (e.g. mutations, small-molecular binding) that can impact flexibility. In future studies, it will be interesting to see the many ways in which flexibility can help orchestrate the dynamics of these assemblies.

With the ever-increasing availability of computational facilities, the ribosome is now becoming a model system for exploring theoretical concepts in biomolecular dynamics. That is, while performing a ribosome simulation used to represent a major technical accomplishment, the field is now entering a stage where the primary challenge is to craft pointed questions, as well as suitable models for addressing them. This stage of development is reminiscent of the protein folding field in the late 1990s and early 2000s. At that time, there was an endless stream of proposed theoretical models, where each could be tested by applying simulation techniques. 20 years later, similar approaches are becoming possible for large assemblies, such as the ribosome. Building on current efforts, we anticipate that the continued integration of experiments, theoretical concepts and simulation techniques will allow for the identification of the precise molecular factors that control complex biomolecular assemblies.

## Author contributions

Conceptualization, P.C.W. and G.F.; methodology, F.C.F., R.J.O., P.C.W.; formal analysis, all authors; writing—original draft preparation, F.C.F. and P.C.W.; writing—review and editing, all authors. All authors have read and agreed to the published version of the manuscript.

## Data availability

All simulated data is available upon reasonable request.

## Acknowledgments

We would like to acknowledge generous support from the Northeastern University Discovery cluster and Northeastern University Research Computing staff.

## Abbreviations

The following abbreviations are used in this manuscript:

rmsf: root mean-squared fluctuations
LSU: large subunit
SSU: small subunit
mfpt: mean first-passage time

## Footnotes

1 See Methods for details.

## References

1. A Korostelev and H F Noller. The ribosome in focus: new structures bring new insights. Trends Biochem. Sci., 32(9):434–41, Sep 2007.

2. Joachim Frank and Ruben L Gonzalez, Jr. Structure and dynamics of a processive brownian motor: The translating ribosome. Annu. Rev. Biochem., 79(1):381–412, 2010.

3. T Martin Schmeing and V Ramakrishnan. What recent ribosome structures have revealed about the mechanism of translation. Nature, 461(7268):1234–1242, 2009.

4. M V Rodnina and W Wintermeyer. The ribosome as a molecular machine: the mechanism of tRNA-mRNA movement in translocation. Biochem. Soc. Trans., 39(2):658–62, Apr 2011.

5. Mikel Valle, Andrey Zavialov, Jayati Sengupta, Urmila Rawat, Mans Ehrenberg, and Joachim Frank. Locking and unlocking of ribosomal motions. Cell, 114(1):123–134, 2003.

6. JA Dunkle, L Wang, M B Feldman, A Pulk, V B Chen, G J Kapral, J Noeske, J S Richardson, S C Blanchard, and J H D Cate. Structures of the bacterial ribosome in classical and hybrid states of tRNA binding. Science, 332(6032):981–4, May 2011.

7. AH Ratje, J Loerke, A Mikolajka, M Brunner, P W Hildebrand, AL Starosta, A Donhöfer, SR Connell, P Fucini, T Mielke, P C Whitford, J N Onuchic, Y Yu, K Y Sanbonmatsu, RK Hartmann, PA Penczek, D N Wilson, and C M T Spahn. Head swivel on the ribosome facilitates translocation by means of intra-subunit tRNA hybrid sites. Nature, 468(7324):713–716, Dec 2010.

8. Srividya Mohan, John Paul Donohue, and Harry F Noller. Molecular mechanics of 30S subunit head rotation. Proc. Natl. Acad. Sci. USA, 111(37):13325–13330, 2014.

9. David J F Ramrath, Hiroshi Yamamoto, Kristian Rother, Daniela Wittek, Markus Pech, Thorsten Mielke, Justus Loerke, Patrick Scheerer, Pavel Ivanov, Yoshika Teraoka, Olga Shpanchenko, Knud H Nierhaus, and Christian M T Spahn. The complex of tmRNA-SmpB and EF-G on translocating ribosomes. Nature, 485(7399):526–529, 2013.

10. Lucas H Horan and Harry F Noller. Intersubunit movement is required for ribosomal translocation. Proc. Natl. Acad. Sci. USA, 104(12):4881–4885, 2007.

11. Peter V Cornish, Dmitri N Ermolenko, David W Staple, Lee Hoang, Robyn P Hickerson, Harry F Noller, and Taekjip Ha. Following movement of the l1 stalk between three functional states in single ribosomes. Proceedings of the National Academy of Sciences, pages pnas-0813180106, 2009.

12. Jingyi Fei, Jonathan E Bronson, Jake M Hofman, Rathi L Srinivas, Chris H Wiggins, and Ruben L Gonzalez. Allosteric collaboration between elongation factor G and the ribosomal L1 stalk directs tRNA movements during translation. Proc. Natl. Acad. Sci. USA, 106(37):15702–15707, 2009.

13. RA Marshall, M Dorywalska, and JD Puglisi. Irreversible chemical steps control intersubunit dynamics during translation. Proc Nat Acad Sci USA, 105(40):15364–9, Oct 2008.

14. Egor Svidritskiy, Axel F. Brilot, Cha San Koh, Nikolaus Grigorieff, and Andrei A. Korostelev. Structures of yeast 80S ribosome-tRNA complexes in the rotated and nonrotated conformations. Structure, 22(8):1210–1218, 2014.

15. Tatyana V. Budkevich, Jan Giesebrecht, Elmar Behrmann, Justus Loerke, David J.F. Ramrath, Thorsten Mielke, Jochen Ismer, Peter W. Hildebrand, Chang Shung Tung, Knud H. Nierhaus, Karissa Y. Sanbonmatsu, and Christian M.T. Spahn. Regulation of the mammalian elongation cycle by subunit rolling: A eukaryotic-specific ribosome rearrangement. Cell, 158(1):121–131, 2014.

16. Yuzuru Itoh, Andreas Naschberger, Narges Mortezaei, Johannes M. Herrmann, and Alexey Amunts. Analysis of translating mitoribosome reveals functional characteristics of translation in mitochondria of fungi. Nat. Commun., 11(1):1–10, 2020.

17. F Tama, Mikel Valle, J Frank, and C L Brooks. Dynamic reorganization of the functionally active ribosome explored by normal mode analysis and cryo-electron microscopy. Proc. Natl. Acad. Sci.USA, 100(16):9319–23, Aug 2003.

18. Y Wang, A J Rader, I Bahar, and R L Jernigan. Global ribosome motions revealed with elastic network model. J. Struct. Biol., 147(3):302–14, Sep 2004.

19. Paul C Whitford, Scott C Blanchard, Jamie H D Cate, and Karissa Y Sanbonmatsu. Connecting the kinetics and energy landscape of tRNA translocation on the ribosome. PLoS Comput. Biol., 9(3):e1003003, 2013.

20. Lars V Bock, Christian Blau, Gunnar F Schröder, Iakov I Davydov, Niels Fischer, Holger Stark, Marina V Rodnina, Andrea C Vaiana, and Helmut Grubmuller. Energy barriers and driving forces in tRNA translocation through the ribosome. Nat. Struct. Mol. Biol., 20:1390–1396, 2013.

21. Lars V Bock, Christian Blau, Andrea C Vaiana, and Helmut Grubmuöller. Dynamic contact network between ribosomal subunits enables rapid large-scale rotation during spontaneous translocation. Nucleic Acid Res., 43(14):6747–6760, 2015.

22. Mariana Levi and Paul Charles Whitford. Dissecting the energetics of subunit rotation in the ribosome. J. Phys. Chem. B, 123:2812–2923, 2019.

23. Paul C Whitford, Jeffrey K Noel, Shachi Gosavi, Alexander Schug, Kevin Y Sanbonmatsu, and Josie N Onuchic. An all-atom structure-based potential for proteins: bridging minimal models with all-atom empirical forcefields. Prot. Struct. Func. Bioinfo., 75(2):430–441, 2009.

24. Jeffrey K. Noel, Mariana Levi, Mohit Raghunathan, Heiko Lammert, Ryan L. Hayes, J N Onuchic, and Paul C. Whitford. SMOG 2: A versatile software package for generating structure-based models. PLoS Comput. Biol., 12(3):e1004794, 03 2016.

25. Paul C Whitford, Wen Jiang, and Philip Serwer. Simulations of Phage T7 Capsid Expansion Reveal the Role of Molecular Sterics on Dynamics. Viruses, 12, 2020.

26. Yong Duan, Chun Wu, Shibasish Chowdhury, Mathew C. Lee, Guoming Xiong, Wei Zhang, Rong Yang, Piotr Cieplak, Ray Luo, Taisung Lee, James Caldwell, Junmei Wang, and Peter Kollman. A point-charge force field for molecular mechanics simulations of proteins based on condensed-phase quantum mechanical calculations. J. Comp. Chem., 24(16):1999–2012, 2003.

27. Jeffrey K Noel, Paul C Whitford, and José N Onuchic. The shadow map: a general contact definition for capturing the dynamics of biomolecular folding and function. J. Phys. Chem. B, 116(29):8692–8702, 2012.

28. Kien Nguyen and Paul C Whitford. Steric interactions lead to collective tilting motion in the ribosome during mRNA-tRNA translocation. Nat. Commun., 7:10586–10586, 2016.

29. E Lindahl, B Hess, and D van der Spoel. GROMACS 3.0: a package for molecular simulation and trajectory analysis. J Mol Mod, 7(8):306–317, 2001.

30. Mark James Abraham, Teemu Murtola, Roland Schulz, Sziliard Piall, Jeremy C. Smith, Berk Hess, and Erik Lindahl. Gromacs: High performance molecular simulations through multi-level parallelism from laptops to supercomputers. SoftwareX, 1-2:19–25, 2015.

31. Huan Yang, Prasad Bandarkar, Ransom Horne, Vitor B.P. Leite, Jorge Chahine, and Paul C. Whitford. Diffusion of tRNA inside the ribosome is position-dependent. J. Chem. Phys., 151(8), 2019.

32. Eric D Hoffer, Samuel Hong, S Sunita, Tatsuya Maehigashi, Jnr Gonzalez, Ruben L, Paul C Whitford, and Christine M Dunham. Structural insights into mrna reading frame regulation by trna modification and slippery codon-anticodon pairing. eLife, 9:e51898, oct 2020.

33. Robert B. Russell and Geoffrey J. Barton. Multiple protein sequence alignment from tertiary structure comparison. Proteins, 14(2):309–323, 1992.

34. W Humphrey, A Dalke, and K Schulten. VMD: Visual molecular dynamics. J. Mol. Graph., 14(1):33–38, FEB 1996.

35. K Okazaki, N Koga, S Takada, J N Onuchic, and P G Wolynes. Multiple-basin energy landscapes for large-amplitude conformational motions of proteins: Structure-based molecular dynamics simulations. Proc Nat Acad Sci USA, 103(32):11844–11849, Jan 2006.

36. C Hyeon and D Thirumalai. Capturing the essence of folding and functions of biomolecules using coarse-grained models. Nat. Commun., 2:487, Jan 2011.

37. Michele Di Pierro, Davit A. Potoyan, Peter G. Wolynes, and José N. Onuchic. Anomalous diffusion, spatial coherence, and viscoelasticity from the energy landscape of human chromosomes. Proc. Natl. Acad. Sci. USA, 115(30):7753–7758, 2018.

38. Hue Sun Chan, Zhuqing Zhang, Stefan Wallin, and Zhirong Liu. Cooperativity, local-nonlocal coupling, and nonnative interactions: principles of protein folding from coarse-grained models. Annu. Rev. Phys. Chem., 62, 2011.

39. Jeffrey K Noel and Paul C Whitford. How EF-Tu can contribute to efficient proofreading of aa-tRNA by the ribosome. Nat. Commun., 7:13314, 2016.

40. Kien Nguyen, Huan Yang, and Paul Charles Whitford. How the ribosomal A-site finger can lead to tRNA species-dependent dynamics. J. Phys. Chem. B, 121:2767–2775, 2017.

41. Mariana Levi, Kelsey Walak, Ailun Wang, Udayan Mohanty, and Paul Charles Whitford. A steric gate controls P/E hybrid-state formation of tRNA on the ribosome. Nat. Commun., 11:5706, 2020.

42. Mariana Levi, Jeffrey K. Noel, and Paul C. Whitford. Studying ribosome dynamics with simplified models. Methods, 162-163:128–140, 2019.

43. Robert B Best and Gerhard Hummer. Reaction coordinates and rates from transition paths. Proceedings of the National Academy of Sciences of the United States of America, 102(19):6732–6737, 2005.

44. P Das, M Moll, H Stamati, LE Kavraki, and C Clementi. Low-dimensional, free-energy landscapes of protein-folding reactions by nonlinear dimensionality reduction. Proc. Natl. Acad. Sci. USA, 103(26):9885–9890, Jan 2006.

45. RB Best, E Paci, G Hummer, and OK Dudko. Pulling direction as a reaction coordinate for the mechanical unfolding of single molecules. J. Phys. Chem. B, 112(19):5968–5976, MAY 15 2008.

46. Sergei V Krivov. On reaction coordinate optimality. J. Chem. Theory Comput., 9(1):135–146, 2013.

47. Dmitrii E. Makarov. Barrier Crossing Dynamics from Single-Molecule Measurements. J. Phys. Chem. B, 125(10):2467–2476, 2021.

48. Robert B Best and Gerhard Hummer. Reaction coordinates and rates from transition paths. Proc. Natl. Acad. Sci. USA, 102(19):6732–6737, 2005.

49. G Hummer. Position-dependent diffusion coefficients and free energies from bayesian analysis of equilibrium and replica molecular dynamics simulations. New J. Phys., 7:34–34, Feb 2005.

50. F Lacroute. RNA and protein elongation rates in Saccharomyces cerevisiae. MGG Molecular & General Genetics, 125(4):319–327, 1973.

51. Jeffrey K Noel, Jorge Chahine, Vitor BP Leite, and Paul Charles Whitford. Capturing transition paths and transition states for conformational rearrangements in the ribosome. Biophys. J., 107(12):2881–2890, 2014.

52. A R Atilgan, S R Durell, R L Jernigan, M C Demirel, O Keskin, and I Bahar. Anisotropy of fluctuation dynamics of proteins with an elastic network model. Biophys. J., 80(1):505–15, Jan 2001.

53. O Miyashita, JN Onuchic, and PG Wolynes. Nonlinear elasticity, proteinquakes, and the energy landscapes of functional transitions in proteins. Proc. Natl. Acad. Sci. USA, 100(22):12570–12575, Jan 2003.

54. Yibing Shan, Anton Arkhipov, Eric T. Kim, Albert C. Pan, and David E. Shawa. Transitions to catalytically inactive conformations in EGFR kinase. Proceedings of the National Academy of Sciences of the United States of America, 110(18):7270–7275, 2013.

55. Jovaun Jackson, Kien Nguyen, and Paul Charles Whitford. Exploring the balance between folding and functional dynamics in proteins and RNA. International Journal of Molecular Sciences, 16(4):6868–6889, 2015.

56. C W Müller, G J Schlauderer, J Reinstein, and G E Schulz. Adenylate kinase motions during catalysis: an energetic counterweight balancing substrate binding. Structure, 4(2):147–56, Feb 1996.

